# Memorability shapes perceived time (and vice versa)

**DOI:** 10.1101/2023.09.02.556045

**Authors:** Alex Ma, Ayana Cameron, Martin Wiener

## Abstract

Visual stimuli are known to vary in their perceived duration. Likewise, some visual stimuli are also known to linger for longer in memory. Yet, whether or not these two features of visual processing are linked is unknown. Despite early assumptions that time is an extracted, or higher-order feature of perception, more recent work over the past two decades has demonstrated that timing may be instantiated within sensory modality circuits. A primary location for many of these studies is the visual system, where duration sensitive responses have been demonstrated. Further, visual stimulus features have been observed to shift perceived duration. These findings suggest that visual circuits mediate or construct perceived time. Here, we present across a series of experiments evidence that perceived time is affected by the image properties of scene size, clutter, and memorability. More specifically, we observe that scene size and memorability dilate time, whereas clutter contracts it. Further, the durations of more memorable images are also perceived more precisely. Conversely, the longer the perceived duration of an image, the more memorable it is. To explain these findings, we applied a recurrent convolutional neural network (rCNN) model of the ventral visual system, in which images are progressively processed over time. We find that more memorable images are processed faster, and that this increase in processing speed predicts both the lengthening and increased precision of perceived durations. These findings thus provide a new avenue in vision research towards the study of perceived image durations as means of explaining visual system responses.

## Introduction

Time is an intrinsic feature of sensory perception. Indeed, all sensory processes must unfold over time. Yet, “time” in itself is a rarely studied feature of perceptual processing. That is, how do we perceive its passage, and how does its passage influence the processing of other features? This presents both a missing aspect of our models of neural functioning and an opportunity for future research: *how is* time *instantiated within sensory processing hierarchies*. Early research on the study of time focused on amodal properties of its perception. That is, with no dedicated sense organ for time, the study of interval timing instead focused on time as a higher order property of perception and cognition (van, 2009).

Within psychology, the dominant model for studying time has been Scalar Expectancy Theory (SET;(Gibbon et al., 1984)), later expanded with the Attentional Gate Model (AGM) of time (Block & Zakay, 1997). Both models assume a pacemaker-accumulator framework, in which clock-unit “ticks” are accumulated until a given threshold. Yet, despite the support of SET and AGM for describing a variety of behavioral features of timing in humans and animals, work conducted throughout the 2000s and 2010s began to reveal perceptual biases that could not be explained by these models. Specifically, *the sensory properties of timed stimuli altered their perceived duration*. Early work in this regard demonstrated that the general magnitude of a stimulus influenced time in a linear manner: “larger” magnitude stimuli, such as size, brightness, loudness, number, numerosity, and speed led to “longer” perceived intervals (i.e. time dilation; (Matthews & Meck, 2016)). A possible explanation for time dilation effects is that these stimuli drew more attention to them as a consequence of their magnitude (Tse et al., 2004), yet this explanation lacks validity in the AGM model, which would predict that such magnitudes would act as a distraction *away* from time, and so should lead to opposite distortions (i.e. time contraction). Explanations for these findings included a generalized “magnitude” system in the brain (Walsh, 2003), with time being just one aspect, and a basic “energy-readout” model in which stimuli that elicited more activity led to longer intervals (Eagleman & Pariyadath, 2009). Yet, further studies revealed findings inconsistent with these accounts, in which time was dilated by other features, such as a visual stimulus’s color, flicker-rate, or spatial frequency, all of which were non-monotonic (Aaen-Stockdale et al., 2011; Bruno & Cicchini, 2016). Further, stimuli of *less* magnitude could be perceived as longer if the context of an experiment was changed (Matthews et al., 2011). Inter-modal effects also existed, such that visual stimuli were generally perceived as briefer than auditory stimuli of the same duration (Allman et al., 2014). “Higher-order” visual stimuli also dilated time, including body motion (i.e. upright human point-light walkers are perceived to last longer than inverted or scrambled walkers; (Wang & Jiang, 2012), emotional content (i.e. emotional faces and frightening images are longer than neutral faces/images; (Lake et al., 2016)), and scenes (i.e. images of scenes are perceived as longer than scrambled scenes; (Varakin et al., 2013)). For these latter stimuli, an important distinction is that it is their specific *content*, not their complexity, that dilates time; indeed, white noise patterns of differing complexity fail to have any impact on perceived duration (Palumbo et al., 2014).

Extending the results from above, two models have attempted to explain how these effects emerge. The first, proposed by (Ahrens & Sahani, 2011), suggested that the brain develops a Bayesian prior for dynamic stimuli that matches the time-varying statistical structure of the environment. That is, natural images (i.e. movies) typically display a 1/f power spectrum in their variation, which allows for predictions of how a typical scene will unfold over time. As such, the brain can exploit this variation to provide a “readout” of time based on sensory change from one moment to the next. A prediction of this model is that dynamic stimuli that contain 1/f spectra will be perceived better (e.g. more precisely) than stimuli that lack this structure. Several experiments demonstrated this effect using moving “cloud” stimuli where the 1/f structure could be controlled. The second model, proposed by (Roseboom et al., 2019), also provided a change-detection account, but without any assumed prior, and instead presumed that the perception of time arose by measuring relative momentary differences between successive video frames. A prediction of this model was that movies containing more changes (e.g. a busy city street) should lead to longer time estimates than those with less change (e.g. an empty field). This prediction was confirmed in experimental data from human observers. The work described above, when applied to the visual system, suggests a hierarchy of time dilation effects. That is, a variety of features from low to high level have been found to influence perceived duration. Yet, the majority of time dilation effects have involved lower-levels of the hierarchy, manipulating simple features such as size, contrast, color, etc. Further, many of these effects have favored stimulus manipulations selective to the dorsal visual stream. Yet, stimuli putatively driven by the ventral stream can also dilate time (Cicchini, 2012), which may be driven by their semantic content, rather than low-level sensory features (Varakin et al., 2013; Suárez-Pinilla et al., 2019). However, previous research using high-level visual images (Cardaci et al., 2009; Varakin et al., 2013; Folta-Schoofs et al., 2014) did not account for semantic properties.

As a means of studying the visual system, one fruitful approach is to employ the use of artificial neural networks (ANNs). ANNs, which may be divided into either recurrent (RNN) or convolutional (CNN) types, have shown great promise over the past decade for modeling neural processes, providing both insights and predictions regarding a variety of sensorimotor phenomena. CNNs in particular have been used both as a means of providing superior image recognition and classification, and as a model of the visual system. For the study of time perception, both RNNs and CNNs have been employed to great effect (Goudar & Buonomano, 2018; Bi & Zhou, 2020). Going forward, we focus here on CNNs due to their link back to the visual system. Notably, only one CNN has been linked to time perception (Roseboom et al., 2019; Sherman et al., 2022; Fountas et al., 2022). Specifically, in the study of Roseboom and colleagues (2019), where movies of city scenes were judged longer than outdoor nature scenes, the authors employed a CNN model (AlexNet) in which, at each layer, the Euclidean distance between activation patterns at successive frames was measured and marked as a “change” if it exceeded an adaptive threshold (T) based on the mean number of previous changes. These values were then regressed against the true durations for each of the videos with a support vector machine so as to provide predicted durations for a given input. After training, the model produced the same biases as human observers.

The application of the CNN to human timing data by Roseboom and colleagues has shown the success of using such an approach. Yet, despite this success, less insight has been gained towards a holistic model of how the visual system encodes time. Indeed, change-detection algorithms have been proposed before for studying time perception (Fraisse, 1984; Poynter & Homa, 1983; Brown, 1995), yet can only accommodate a limited number of findings (Liverence & Scholl, 2012; Bruno et al., 2012). The work here therefore seeks to further examine the nature of timing effects on the visual system, with a particular emphasis on ventral stream selective processes.

## Results

### Perceived Time is Influenced by Scene Size and Clutter in Opposite Directions

To begin, we tested an initial group of human subjects (n=52) on a temporal categorization task (Figure 1A), in which they were presented with images for a set of six possible durations on a given trial (log-spaced, between 300 and 900ms). For each image, subjects were required to classify the presented image into “long” and “short” duration categories via a button-press as quickly yet as accurately as possible. We gave subjects no instructions regarding the images themselves, asking them only to attend to the durations they were presented. For this experiment, the images we used were drawn from the Size/Clutter database built and described by (Park et al., 2015) (see Methods). These images represent a series of scenes with normed responses across participants for ratings of scene size or clutter. By example, a scene with a small size but high clutter may be a full pantry, whereas a large size but low clutter scene may be an empty warehouse. The scenes were presented across six levels of size and clutter, for a total of 36 possible combinations (Figure 1B). The data were analyzed via a generalized linear mixed model approach in which the scene size and clutter levels, along with presented duration, were fixed effects and subject was a random effect. Here, we observed an effect of both scene size and clutter, such that models with these terms outperformed models without them [Scene Size: *χ*^2^=99.37, *p<*0.001; Clutter: *χ*^2^=5.94, *p*=0.015]. Strikingly, the direction of each effect moved in opposite directions; Scene size led subjects to categorize stimuli as “long” more often [*β*=0.055, 95% CI: 0.044 — 0.065], whereas clutter led subjects to categorize stimuli as “short” more often [*β*=-0.044, 95% CI: -0.079 — -0.008] (Figure 1B). Additionally, we observed an interaction between presented duration and clutter [*χ*^2^=4.772, p=0.029]. Notably, the slope of this interaction was positive [*β*=0.067, 95% CI: 0.006 — 0.127], such that the slope of the psychometric function was higher for larger levels of clutter (Moscatelli et al., 2012); thus, despite the bias to classify the duration of more cluttered images as “short”, subjects are more precise in their classifications. No such interaction was observed for scene size.

**Figure 1.**
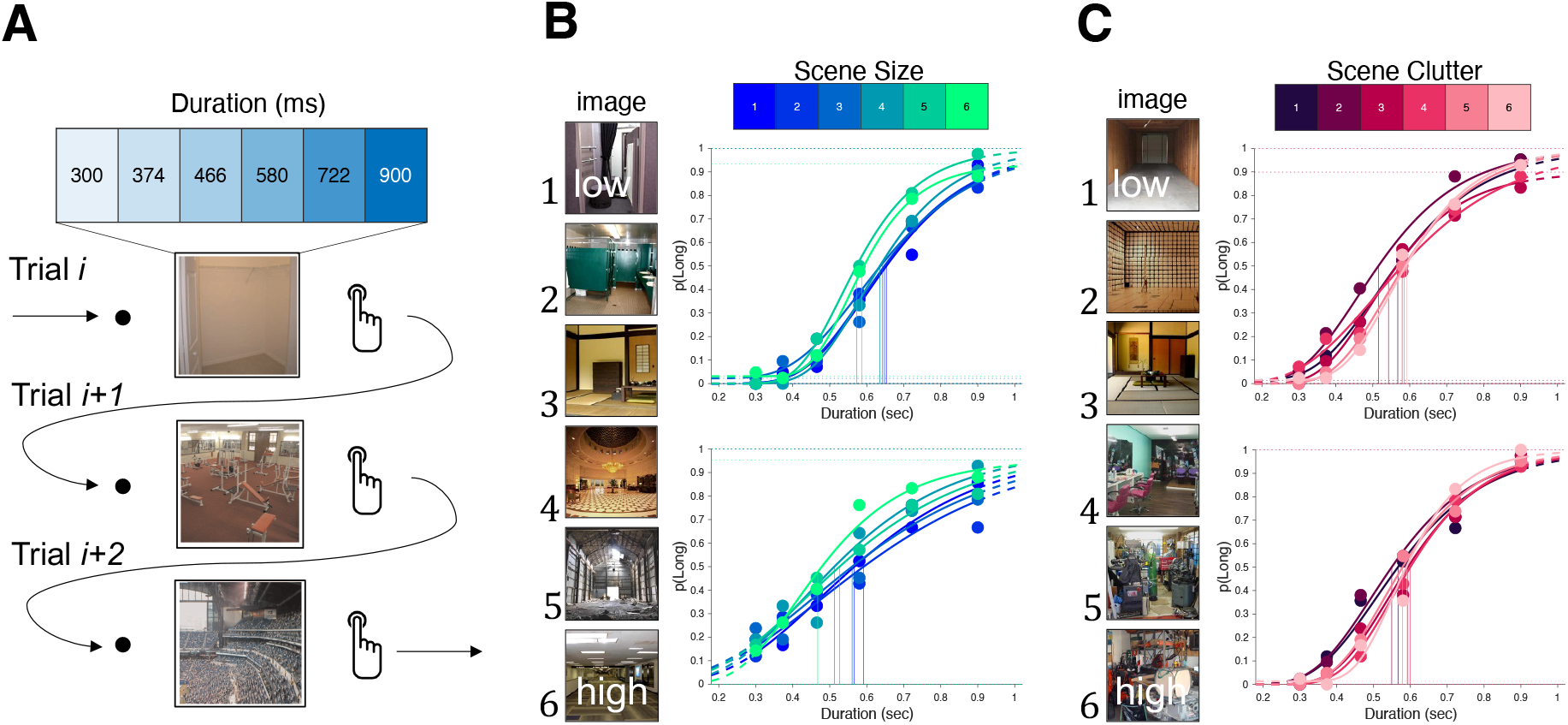
Scene information shifts perceived time. **A**) Schematic for the temporal categorization task; on a given trial (i) subjects viewed a fixation point followed by an image for one of six possible durations between 300 and 900ms. After the image disappeared, subjects were required to classify the image duration as “long” or “short” as quickly yet as accurately as possible, after which the next trial (i+1) began immediately. **B**) Scene size was varied across six levels and was observed to dilate perceived time, such that subjects were more likely to categorize larger scene size images as “long”. Example psychometric functions are presented for two subjects from Experiment 1 (top) and 2 (bottom). **C**) Scene clutter was also varied across six levels and was observed to contract perceived time, such that subjects were less likely to categorize more cluttered images as “long”. Example psychometric curves from two subjects are again presented for Experiments 1 (top) and 2 (bottom). Curves were fit using the psignifit 4.0 toolbox and are presented here for visualization purposes only.

The results of the first experiment consequently showed that scene size and clutter could push perceived duration into two separate directions. We note that this finding goes against a simple attentional explanation, unless one were to suggest a more complicated explanation that scene sizes draw more attention than scene clutter, which decrease attention with greater clutter. Likewise, a magnitude-based effect cannot explain these findings, as both scene size and clutter are larger magnitudes. To further validate these effects, we collected a replication dataset in a new group of subjects (n=50). As an additional control, the images presented were set to grayscale and normalized for luminance (see Methods), to ensure the results were not due to low-level differences in the intensity of the image. Once again, we observed a significant effect of including both scene size [*χ*^2^=9.497, p=0.002] and clutter [*χ*^2^=8.6, p=0.003] in our model, with scene size pushing stimuli to be classified as “long” more often [*β*=0.017, 95% CI: 0.005 — 0.028] and clutter pushing stimuli to be classified as “short” more often [*β*=-0.018, 95% CI: -0.029 — -0.006]. However, a model including an interaction between duration and clutter did not significantly improve the fit [*χ*^2^=0.047, p=0.826], thus failing to replicate the effect of clutter on precision.

### Memorability Lengthens Perceived Time

The results of the first two experiments demonstrated that semantic details of scenes can shift perceived time in different directions, depending on the type of information conveyed. These findings could not be explained by simple magnitude or attention-based theories, nor by differences in low-level features of the images. So, why do these images affect time in different ways? We return to this question in the discussion, but note that the richness of scene images provides a number of distinct cues, many of which are perceived immediately. Beyond features such as size or clutter, an additional feature of images is their intrinsic memorability, or the probability that they will be recalled later (Khosla et al., 2015; Rust & Mehrpour, 2020). Numerous studies have investigated features that give rise to memorability, noting that it is a unique property of images that operates independent of attention (Rust & Mehrpour, 2020). One possibility, then, is that memorability affects perceived time. To explore this possibility, we conducted a third experiment on a new set of subjects, in which subjects categorized the duration of images that varied according to their memorability ratings. Images were uniformly drawn from the Large-Scale Image Memorability dataset (LaMem) across all memorability scores and divided into seven equally-spaced bins from high (1) to low (7) in memorability ratings (Figure 2A).

**Figure 2.**
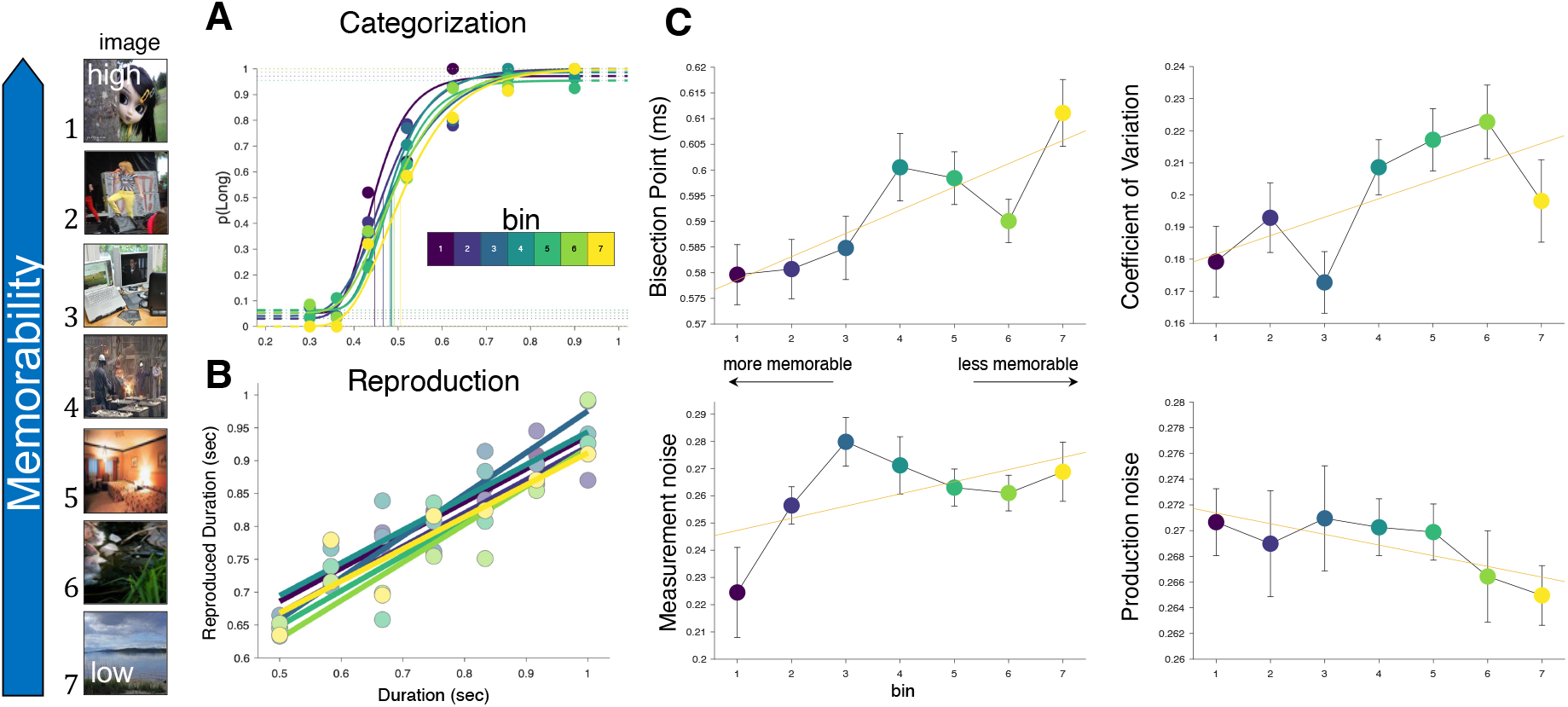
Memorability dilates perceived time. Subjects were presented with stimuli drawn from the LaMem dataset which varied by their memorability ratings and divided into a set of 7 bins from low (7)to high (1). **A**) In a temporal categorization task, subjects were more likely to categorize images with higher memorability scores into the “long” duration category. Panel A displays psychometric functions from an example subject; Panel **C**) left displays average bisection points across the seven memorability bins. Additionally, subjects were more precise at categorizing the durations of higher memorability ratings, as evidenced by reduced coefficient of variation values (Panel C top right). **B**) In a temporal reproduction task (separate subjects), subjects reproduced longer durations after having encoded higher memorability images; data are shown from an example subject. **C**) Bottom panels display the measurement (left) and production (right) noise as derived from a Bayesian Observer model fit to subject responses, in which measurement noise is additionally shown to be reduced for higher memorability images.

Psychometric functions were constructed for the responses proportions for each tested duration, from which the bisection point (BP), defined as the duration at which subjects were equally likely to classify the interval as “long” or “short”, and the coefficient of variation (CV), defined as half the difference between upper and lower thresholds divided by the BP, were calculated (see Methods). A repeated-measures ANOVA of BP values found a significant effect of memorability [*F* (6,150)=3.467, *p*=0.003, *η*^2^_p_=0.122], which was observed to be linear in nature such that subjects were more likely to classify intervals as “long” for more memorable images [*t* (150)=3.827, *p<*0.001] (Figure 2A,B). Surprisingly, for the CV, we also detected a significant effect of memorability [*F* (3.738,97.99) = 2.653, *p* = 0.041, η^2^_p_ = 0.093] that was also linear in nature, such that more memorable images were also classified with *better* precision [*t* (156)=2.643, *p=*0.009] (Figure 2C).

### Perceived Time Increases Memorability

The results of Experiment 3 demonstrated, strikingly, that more memorable images are both perceived as longer than less memorable ones and more precisely. That is, an intrinsic aspect of these images that allows them to be better recalled is also responsible for dilating the duration which they are presented for. Yet, this relationship is correlational in nature, and so the directionality of the effect is unknown (Figure 3A). To pose the question clearly: do these images last longer because they are more memorable, or are they more memorable because they last longer? Indeed, previous research has shown that the duration for which an image is *objectively* presented increases the likelihood that it will be remembered (Potter & Levy, 1969; Potter, 2012; Wichmann et al., 2002), yet whether a *subjectively* longer image is thus recalled better is not known. Evidence of such a relationship would differ from a magnitude-based explanation; for example, larger stimuli are commonly perceived as lasting for a longer duration, but presenting a stimulus for a longer duration does not make it appear larger.

**Figure 3.**
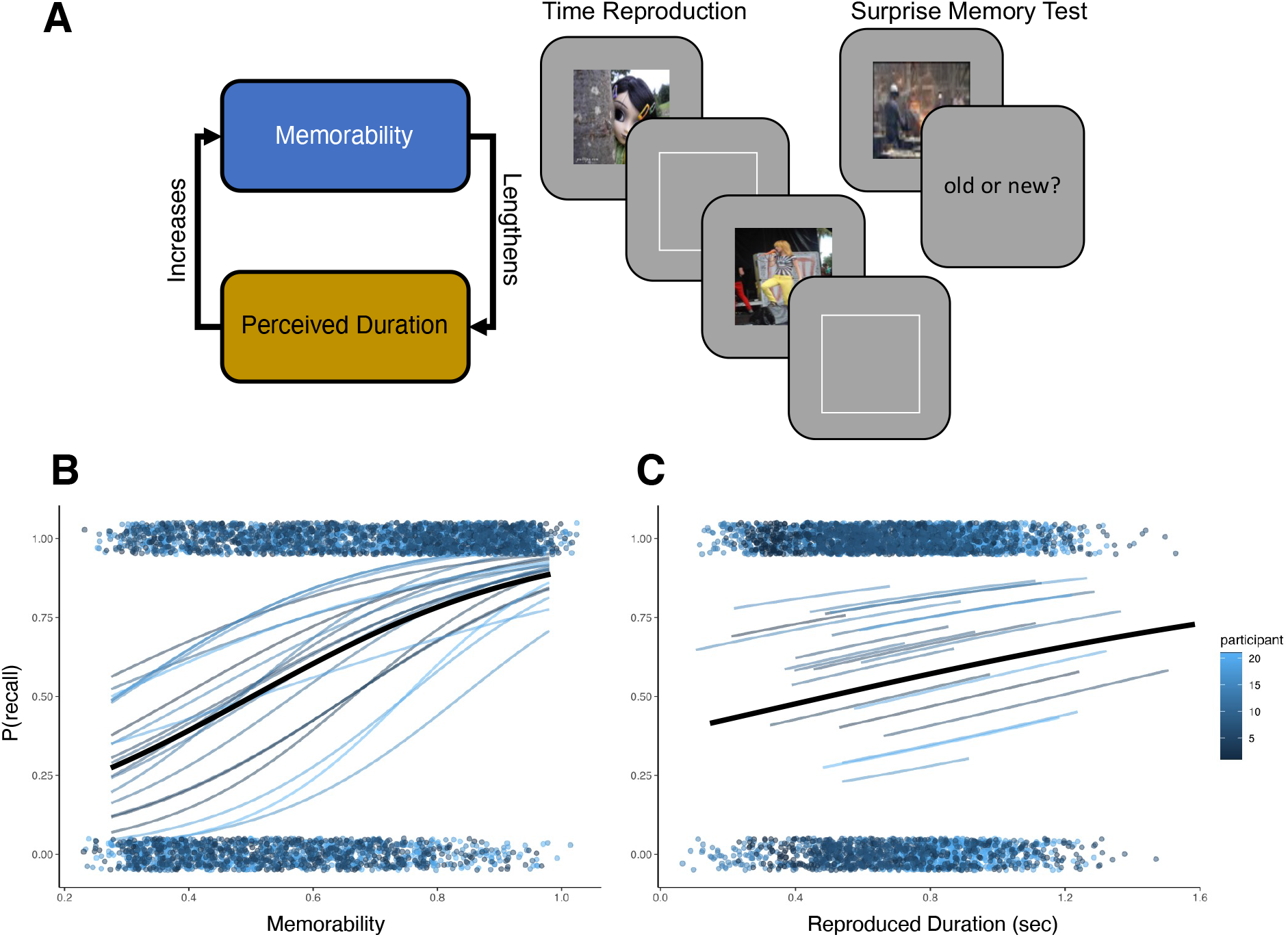
Perceived duration affects memorability. **A**) Proposed bidirectional relationship between memorability and perceived duration, such that more memorable stimuli increase the length of perceived durations, but longer durations also increase the likelihood of remembering a stimulus. To test this, subjects performed a time reproduction task with memorability stimuli, and then performed a surprise memory test on a subsequent day in which they recalled stimuli from the previous day. **B**) Regression estimates for single subjects and group average (black line) between memorability of presented images and recall performance demonstrating greater probability of recall for more memorable images. **C**) Regression estimates for average reproduced duration estimates for individual images and recall performance also demonstrating greater probability of recall for longer reproduced durations.

To test this hypothesis, we had a new set of subjects perform a temporal reproduction task using the same memorability images from Experiment 3 (Figure 3A). In this task, subjects were presented with images from the memorability image set for a random interval between 500-1000ms, and then required to reproduce that interval by pressing and holding a response key for the same interval. We chose a reproduction task here for two reasons: 1) to replicate the findings of Experiment 3 but in a different task, and 2) to obtain a continuous, rather than categorical, estimate of perceived duration. The result of this initial task replicated the findings of experiment 3; a linear mixed model of reproduced durations was significantly improved by adding the memorability score of the image [*F* =9.567, *p*=0.002], with higher memorability scores associated with longer duration estimates [*β*=0.032, 95% CI: 0.011 — 0.052]. As an additional measure, we decomposed reproduced duration estimates with a Bayesian observer model, in which the measurement of durations on each trial are conceived as draws from a noisy Gaussian distribution that scales with the interval duration. These estimates are then combined optimally with a uniform prior distribution of presented durations to form a posterior estimate, which is then further corrupted by motor production noise in the reproduction phase (Jazayeri & Shadlen, 2010; Remington et al., 2018; De et al., 2021; De et al., 2023). Fitting this model to single-trial responses yields an estimate of both the measurement and production noise widths. Here, we observed that the measurement noise decreased for images from higher memorability bins [*F* (3.804,68.476)=2.611, p=0.045, η^2^_p_=0.127] in a linear manner [*t* (108)=2.163, *p*=0.033], while no effect was found for production noise [*F* (4.728, 89.829)=0.465, p=0.791]. Thus, a similar effect of memorability on the CV of Experiment 3 was also observed for the measurement error of Experiment 4.

Following the reproduction tasks, all subjects returned a day later for a second session, in which they were presented with a surprise memory test (Figure 3A). In this phase, subjects were presented with the same 196 images from the previous day, along with a new set of 196 image foils drawn from the same memorability bins as the first set. Subjects were presented with each image and asked to judge if they had seen them on the previous day. A generalized linear mixed model analysis of accuracy scores in this task for each image replicated the well-known effect of memorability [*χ*^2^=72.613, p*<*0.001], with higher memorability scores associated with a greater probability of recall [*β*=5.223, 95% CI: 4.661 — 5.783]. Crucially, the inclusion of average reproduced duration from the previous day’s session also improved model fit [*χ*^2^=4.506, p=0.033], with longer reproduced durations associated with greater recall [*β*=0.698, 95% CI: 0.061 — 1.335]. We note that the intervals used represented the average across all objectively presented durations for each image (see Methods). An interaction between memorability and reproduced duration did not significantly improve the fit and so was not warranted [*χ*^2^=0.07, p=0.791] (Figure 3B). Inspection of predicted model fits additionally yielded an unexpected finding: while longer duration estimates were associated with better recall, those subjects who overall reproduced longer durations were *less* likely to recall images in general. This finding, an example of Simpson’s paradox, was evident when removing subject as a random effect, which thus changed the beta estimate for duration from a positive value to a negative one [*β*=-0.698, 95% CI: -0.061 — -1.335]. One possible explanation for this effect is that longer duration estimates are typically associated with greater attention to time. It is possible, therefore, that subjects who were able to more effectively able to ignore the images, and so reproduce longer duration estimates, were thus less effective at encoding the images into memory; nonetheless, these same subjects were still affected by the intrinsic memorability of those images, such that more memorable ones, and those reproduced as relatively longer, were relatively better remembered.

### Neural Network Modeling

How to explain the effect of memorability on time? We assert that appeals to other perceptual phenomena such as attention or magnitude are insufficient to explain this link. Memorability draws on a variety of details that give rise to its effect; further, memorable images exist independent of attentional effects (Bainbridge, 2020; Wakeland-Hart et al., 2022), and one would not judge that a more memorable image is higher along an axis of magnitude like size or quantity. To explain these findings, we turned to computational models of vision. In particular, convolutional neural network (CNN) models have been a major tool for vision researchers to measure corresponding links between activation “layers” and corresponding components of the ventral visual stream. Indeed, CNNs and other model developments (i.e. ResNets) have been quite successful at estimating memorability and linking them to particular image features at a variety of levels of the hierarchy. However, CNNs as a base model cannot provide a strong explanation for our findings, as these models do not operate over time. A recent advance in CNN models however is to add feedback connections within and between layers, thus generating *recurrent* CNN models (rCNN). In these models, the outputs of individual layers can be “unrolled” across successive timesteps as recurrent, feedforward and feedback connections (van & Kriegeskorte, 2020). This process thus provides a timescale by which an input can be successively processed. We note however that the step distance in this case is arbitary; that is, one can appeal to conduction delays between layers, but without neural data to corroborate the difference between so-called “engineering” time and “biological” time is irrelevant (Spoerer et al., 2020).

To investigate how computational models of vision might explain our findings, we turned to a rCNN model known as *BLnet* (Bottom-up Lateral network), containing seven layers and recurrent connections within each layer. We chose this network first because it provides a built-in series of time steps (8) for processing an image, in which a readout is provided with each time step, but more importantly because the output of this network has been shown to correlate with human reaction times for image classification (Spoerer et al., 2020), as well as rapid object recognition (Sörensen et al., 2023). This is achieved by extracting the softmax readout at each of the eight timesteps and then calculating the entropy of each readout. As the model will converge on a set of image categories over others with repeated recurrent steps, the entropy of the softmax distribution will decrease with successive timesteps. By selecting a threshold for entropy, one can infer the model’s “reaction time” to a particular stimulus.

**Figure 4.**
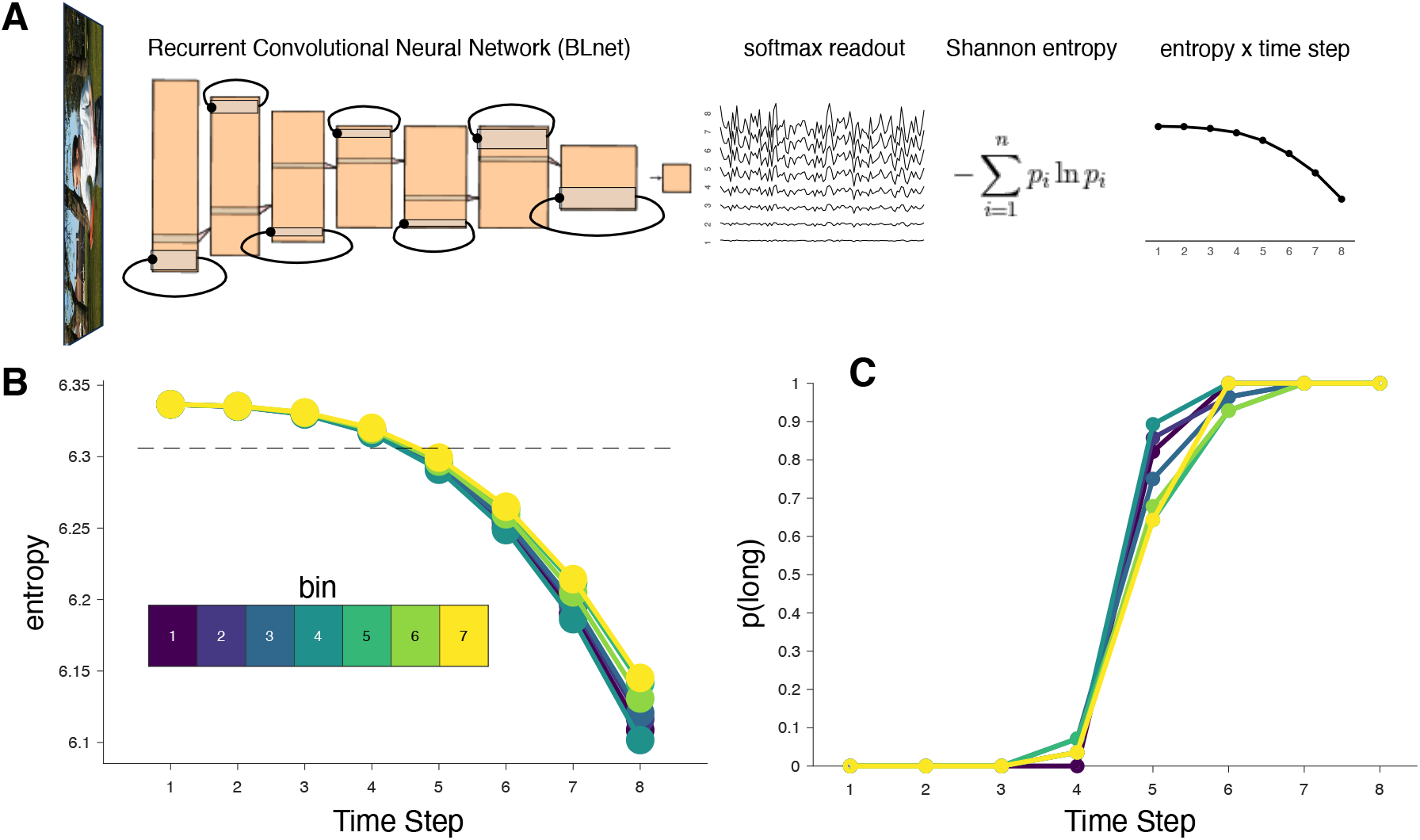
Neural network modeling of memorability and time. **A**) Schematic layers used for a recurrent convolutional neural network (rCNN), *BLnet*. Images are fed and processed across the network in a feedforward manner but with lateral recurrent steps for each layer, such that the input to each layer is refined at successive steps. At the final layer, a softmax readout provides classification probabilities for each timestep (eight timesteps used). For each readout, Shannon entropy is calculated, resulting in a timeseries of entropy values. **B**) The Memorability images used in Experiments 3 and 4 were fed into BLnet, from which entropy values were averaged across each memorability bin, thus revealing that images with higher memorabilities exhibited faster reductions in entropy. Setting a threshold on these entropy values allowed us to construct a psychometric function for categorizing time intervals (for arbitrary time steps) as short or long. The dashed line represents the entropy threshold after fitting the model to human participant choices. **C**) Resulting psychometric functions for the entropy threshold and for each memorability bin. The psychometric data recapitulated both the time dilation effect for more memorable images, as well as the increase in precision.

To begin, we fed the 196 images presented to subjects in Experiments 3 and 4 into *BLnet* and calculated the entropy across the eight timesteps. These responses were then binned by their memorability ratings. As in previous studies, we observed that the entropy decreased across timesteps; notably, we observed that this decrease was logarithmic in nature. Strikingly, we observed that memorability affected the rate of this decrease, such that more memorable images decreased at a faster rate then less memorable ones [timestep x memorability interaction *F* (1,194)=4.487, *p=*0.035] An interpretation of this finding is that images that are more memorable are processed faster than those that are less memorable, with the network converging on a set of categorizations more consistently over time. One possibility, then, is that longer perceived durations are the result of this faster speed with which the network operates are more memorable images. To determine if this was the case, we set an entropy threshold and categorized, for each timestep and each image, that an entropy value above this threshold would categorize the stimulus as “short” and a value below it would be categorized as “long”. The average proportion of “long” responses was then calculated for each memorability bin, thus providing seven psychometric functions which were fit in the same manner as Experiment 3 (see Methods). We then fit these functions to the subject data by finding the entropy threshold which provided the best match for both the bias and precision effects we observed. Remarkably, we found that the model recapitulated *both* of the observed effects in humans, with higher memorability images associated with a greater probability of categorizing a stimulus as “long” and also a steeper -more precise -psychometric function. To the former effect, longer perceived durations were a direct result of the faster speed with which the network converged on a solution, such that the entropy threshold was hit earlier in time. To the latter effect, the increase in precision was also due to the speed at which the network converged, such that the a smaller proportion of the range of possible entropy values for a more memorable images overlapped with the entropy threshold, an observation that we note is similar to accumulator-based models of time perception (Allman et al., 2014).

## Discussion

The results of the preceding experiments demonstrate that higher-order semantic features of scenes can shape perceived time. These features include aspects of scenes relevant for navigation, including scene clutter and size, as well as the intrinsic feature of image memorability. Further, the results show that this effect is bidirectional, such that changes in perceived time have relevance to the perception of those stimuli themselves, such that subjectively longer perceived intervals are more likely to be recalled. These effects point to a series of loci along the ventral visual stream for compressing and dilating subjective time. Combined with our application of a model of the visual system, these effects suggest that the source of changes in perceived time are the result of processing efficiency for those natural images, which occurs as a result of recurrent, feedback connections within visual circuits.

### Scene Features and Perceived Time

The first two experiments conducted demonstrate opposite directions for the image scene qualities of size and clutter on perceived time, such that the former dilates and the latter compresses perceived time. Notably, these findings stand in opposition to a number of explanations for time dilation phenomena. For example, magnitude accounts would predict that both scene size and clutter should produce time dilation effects. Indeed, the perception of clutter is akin to numerosity, where time dilation effects have been observed in response to large dot arrays (Bueti & Walsh, 2009). Likewise, appeals to increases in arousal, attention, or neural response to the stimuli would all suggest that both scene size and clutter should increase perceived duration (Matthews & Meck, 2016). The most similar finding to the present one is a brief report noting that paintings of increasing complexity, defined using low-level features such as edges and contrast, also lead to time compression (Cardaci et al., 2009); yet this study presented images for very long intervals (∼30-60s), whereas the present study employed very brief durations all less than one second. One possibility may relate to the stability of these images across the visual hierarchy. Indeed, while scene clutter is known to peak earlier in decodability of neural responses than scene size, it has also been shown that increases in clutter impair object recognition (Manassi & Whitney, 2018), which may relate to how consistent these representations are in visual responses (Martin et al., 2017; Graumann et al., 2022; Park et al., 2015). Recent work has also shown that increases in the objective time of presented images leads to more stable representations, rather than extended firing rates, in higher-level parts of the ventral stream (Vishne et al., 2023).

A second possibility for explaining the size/clutter effects relates to their actionability. That is, the clutter or size of a scene are both relevant features for navigation (Learmonth et al., 2002), and recent work has shown that humans extract information about a presented scene in a way that supports their movement through that space (Bonner & Epstein, 2017). Indeed, prior work has also demonstrated differentiated neural representations for scenes depending on whether the objects presented in that scene are reachable or not (Josephs & Konkle, 2020). Increases in the size of a scene would suggest a longer necessary path to traverse the space presented, and previous work has shown that larger presented distances dilate perceived time (Riemer et al., 2018). Similarly, a more cluttered scene would suggest more difficulty in reaching ones goal. Yet, this would predict an interaction between scene size and clutter, which was not observed.

### Memorability and Perceived Time

The last two experiments demonstrated that the memorability of a scene dilates the perceived duration for which it was presented. Likewise, increases in the perceived duration of a scene also increase its memorability. These findings go beyond a simple unidirectional explanation, in which the perceptual feature affects its perceived duration as a byproduct of its processing. Indeed, memorability has been demonstrated as a perceptual feature of a stimulus related to processing in higher regions of the ventral stream such as the inferotemporal (IT) cortex (Rust & Mehrpour, 2020; Jaegle et al., 2019). Similar to our findings, recent work has shown that increasing either the objective or *subjective* size of an image also increases its memorability (Masarwa et al., 2022; Jeong, 2023). These findings are suggested to relate to the size and spread of activation resulting from larger images on the surface of striate and extrastriate regions (Pooresmaeili et al., 2013). Comparatively, longer objective durations are also known to increase the memorability of an image (Potter & Levy, 1969; Potter, 2012; Wichmann et al., 2002). Yet, in our findings, we also observed that the durations of more memorable images were perceived with greater precision. That is, a more memorable image is both perceived as longer and more consistent. This second finding sets memorability effects apart from other time dilation effects, which commonly do not change precision, but also points to differences in the stability of the neural representation.

The finding that memorability and time both affect one another suggests a single underlying factor driving both effects. Yet, more generally, why should time dilation effects occur at all? Up until now, the dominant framework asserts that time dilation results from increases in attention, or population neuronal responses. Either case relies on a *more is more* connection, where increases in attention or firing rate lead to longer perceived durations. Crucially, there is a directional link to this framework, where time dilation is a *consequence* of increased neuronal firing, rather than a *cause*. The result is that time dilation is an epiphenomenon operating downstream from neuronal computations for stimulus coding (Noguchi & Kakigi, 2006). However, an alternative framework that we propose here is that time dilation effects may instead serve a *purpose* for the visual system. Under this new framework, we propose that time in the brain serves as an *information seeking strategy (Matthews & Meck, 2016)*. This new framework connects to the recent notion of *priority coding* in the visual system (Rust & Cohen, 2022), wherein stimuli are processed according to the priority they engender (e.g., threatening/emotional/rewarding/appetitive/etc.). Yet, under the priority coding framework, the processing of visual stimuli is limited by an information bottleneck that is time-limited (i.e. only so much information can be processed at once; (Rust & Palmer, 2021)). To surmount this, we suggest that the information bottleneck can be dynamically changed to accommodate higher priority stimuli. In this way, *time is dilated or compressed in order to increase the amount of information that can be processed in any given instance*, and so time is not epiphenomenal, but central to population coding. We note that versions of an adaptive temporal window have been proposed before (e.g. (White, 2017; Pereira et al., 2022)), yet not comprehensively explored across the visual hierarchy.

The notion of changes in information processing related to time has other experimental evidence to support it. For example, humans are able to adjust the rate of evidence accumulation to the rate of stimulus presentations in a dynamic environment (Ossmy et al., 2013). Likewise, recent work has shown that humans can vary the encoding speed of visual items in working memory, depending on the duration at which items are presented (de et al., 2023). These findings provide further support to the notion that time is a controllable feature of visual processing. In support of this, our application of a rCNN model provides an avenue by which time dilation effects may occur. In this model, the responses at each layer are both fed forward to subsequent layers and laterally back to themselves. This process, in which a CNN is “unrolled” in time, allows for image processing to occur across multiple timesteps (Spoerer et al., 2020; Kietzmann et al., 2019), which are meant to mimic both the feedforward “sweep” of the visual hierarchy as well as recurrent connections (Lamme & Roelfsema, 2000; van & Kriegeskorte, 2020). Critically, for image processing, this allows image representations to be refined over time as the model converges on a set of solutions (i.e. the probability distribution of object identities). Here, we observed that more memorable images were processed *faster* across successive timesteps in the rCNN, such that the probability distribution converged earlier in time. By applying a threshold to the timeseries and using this to mimic the decision process in a categorization task, we found that the model could replicate both the time dilation and precision effects observed in behavior for memorability. Time dilation thus results from a faster *speed* of the network, rather than an increase in neural firing; likewise, faster speeds are associated with less variability, and so this leads to greater precision for categorizing time intervals. This finding mirrors recent neural recordings in nonhuman primates, as well as modeling with with RNNs, demonstrating that perceived duration is the result of changes in the speed of neural trajectories through state space (Goudar & Buonomano, 2018; Bi & Zhou, 2020; Wang et al., 2018). In our case, the change in speed is the result of changes in processing *across* layers of the ventral stream, rather than within any given region, a finding that will need to be tested experimentally.

## Conclusion

The results of the experiments outlined here provide evidence for a link between the perception of time and the semantic features of scenes. Further, they indicate a bidirectional effect between memorability and perceived duration. These results point to a framework in which time dilation is both the result and the cause of priority coding in the visual system, which is verified by computational modeling of the ventral visual stream. We suggest that a large variety of visual stimuli may be used to explore timing responses across different levels of the hierarchy, including those associated with reachable objects, animacy, sizes, textures, metamers, and forms, all of which can provide insight to the location of time dilation effects across the visual system.

## Methods

### Participants

A total of 170 participants took part in the four experiments described in this study. No subjects participated in more than one experiment. All subjects were drawn from the undergraduate pool of George Mason University and took part for course credit. All subjects provided informed consent and all procedures were approved by the Institutional Review Board at George Mason University. Experiments 1-3 were run online during the Covid-19 pandemic, whereas Experiment 4 was run in-person after pandemic protocols had been eased for data collection. All subjects were right-handed and neurologically healthy with normal or corrected-to-normal vision. Experiment 1 included 52 subjects, Experiment 2 included 50 subjects, Experiment 3 included 48 subjects, and Experiment 4 included 21 subjects. All experiments were programmed using Psychopy (www.psychopy.org). Online experiments were conducted using the Pavlovia platform (www.pavlovia.org). In-person experiments were conducted in a testing room with stimuli presented on a 100Hz Dell Gaming Monitor and responses collected on a Corsair MX Gaming Keyboard with a 1000Hz polling rate. For all experiments below, subjects were not informed to the nature of the images they were presented, and were not given any instructions related to their processing. Rather, subjects were only told to attend to the duration for which they were presented, regardless of their content.

### Experiments 1 & 2: Temporal Categorization of Scene Size and Clutter

All subjects performed a visual temporal categorization task (also referred to as a time bisection task) with sub-second stimuli. The stimuli consisted of images drawn from the Size/Clutter database of (Park et al., 2015), which is available at https://konklab.fas.harvard.edu/#. A total of 252 images were chosen from across the dataset, which spans six levels of “size” and “clutter”, based on participant ratings. For Experiment 1, we used the images as provided from the Size/Clutter database; for Experiment 2, all images were processed via the SHINE toolbox (Willenbockel et al., 2010), in which the images were turned to grayscale and normalized for luminance. At the start of each trial, participants were presented with a fixation point that appeared at the center of the screen for 500 ms before immediately presenting a visual stimulus. The stimuli order was randomized for each trial and images appeared for one of six logarithmically-spaced time intervals ranging from 300 to 900 milliseconds. Logarithmic spacing allows for more supra-geometric spread when visualizing the data (Kopec & Brody, 2010). Accordingly, each image was presented once for each of the six possible durations, leading to a total of 1512 trials in a given session. A break was implemented every 168 trials, which subjects could end by pressing a response key. The image size was set to (0.5)^2^ height of the monitor, following recommendations for presenting stimuli for online experiments to account for differing screen sizes across subjects. On a given trial, participants were tasked with judging whether the stimulus presented was “short” or “long” based on their subjective threshold of the durations. They were directed to respond as quickly and accurately as possible using the “s” key for “short” and the “l” key for “long”. There was no response screen following the stimulus, participants were simply instructed to answer as soon as the image disappeared. They did not receive feedback during this task and the next trial began upon their response.

#### Analysis

Subject responses were entered in a generalized linear mixed model (GLMM), with stimulus duration and the magnitude of the size or clutter of each image as fixed effects and subject as a random effect. Trials were filtered based on reaction times; we set limits for trialwise RTs to be above 100ms and below 1s. We chose this threshold, rather than a distributional one, to reflect the potentially wider range of RTs resulting from collecting data online. For the GLMM analysis, model comparisons were carried out via Chi Square tests of model complexity. Fixed effects were measured using the likelihood ratio tests.

### Experiment 3: Temporal Categorization of Memorability Images

All subjects performed the temporal categorization task as described for Experiment 1, but with a different set of images. Specifically, we drew a set of images from the Large-Scale Image Memorability Dataset (LaMem; http://memorability. csail.mit.edu/index.html). This dataset contains 60,000 images from a number of distinct sources, each with a corresponding memorability score, reflecting the probability that the image will be recalled later (Khosla et al., 2015). 28 images were randomly sampled from each of seven equally spaced memorability ‘bins’, or ranges of memorability scores [Δbin ∼ 0.10390; Bin 1 = 1 -.89610, Bin 2 = .89610 -.79220, Bin3 = .79220 -.68831, Bin 4 = .68831 -.58441, Bin 5 = .58441 -.48051, Bin 6 = .48051 -.37662, Bin 7 = .37662 -.27273]. Within each bin there was uniform spacing among the scores, ensuring an overall spread across the selected images. The memorability scores were taken from the 2nd training step provided within the LaMem files. This resulted in 196 visual stimuli split among seven different ranges of memorabilities. Additionally, we included here seven possible durations, again log-spaced between 300 and 900ms; this was done to allow for better characterization of the psychometric function for use with fitting routines described below. The combination of seven different memorability ranges and seven possible durations created a total of 42 possible conditions across 196 trials. Every image was seen at all seven durations, resulting in a total of 1372 trials, which was divided into seven blocks to allow participants a break. Thus, each block was about six minutes making the full experiment ∼45 minutes.

#### Analysis

Psychometric functions for each memorability bin were fit using psignifit 4.0 (Schütt et al., 2016). All data were fit using a right-tailed Gumbel distribution to account for the log-spaced nature of the tested intervals, from which the bisection point (BP) and coefficient of variation (CV) were calculated. The BP was determined as the 0.5 point on the curve for categorizing stimuli as long, whereas the CV was defined as half the difference between 0.75 and 0.25 points on the function divided by the BP. As an additional step, we removed any subjects with a BP value that exceeded the tested intervals in the stimulus set or a CV greater than 0.5. Using this conservative threshold, 24 subjects were removed from the analysis.

### Experiment 4: Temporal Reproduction and Recall of Memorability Images

Experiment 4 took place on two separate yet subsequent days. In the first part, subjects performed a duration reproduction task, in which subjects were shown an image and asked to press and hold a button for the same duration of time for which the image was shown (Mioni et al., 2014). The same 196 images from the Experiment 3 were used again for this task. The images were each presented for one of seven possible durations linearly-spaced from 500 to 1000ms such that each duration was represented four times in each bin. On a given trial, subjects were first shown a fixation cross for 500ms, then the image for its specified duration, then asked to reproduce the duration by pressing and holding a response key to match the presented duration. While holding the button down, participants were shown an unfilled white square of the same size as the images as an aid for reproducing the duration. All 196 image reproduction events shown to a participant represented 1 block, and each participant was asked to complete 6 blocks of duration trials, with breaks in between, to finish the first part of the study. Prior to completing 6 blocks of duration trials, participants were asked to complete 3 practice trials, which were equivalent to a typical duration reproduction trial, but with a white unfilled square of the same size as the images. After each practice trial, participants were shown the numerical duration which they reproduced, and the target duration. When they were finished with 3 practice trials, they were asked to complete the normal trials. Each block of 196 trials took about 10 minutes resulting in a total experiment time of about 60 minutes.

The second part was conducted in the same room, using the same monitor and keyboard configuration as the first task. Here, subjects performed a surprise memory recall task, in which they were shown images and asked whether or not they were shown in the duration reproduction task. Subjects were not informed that they would perform the memory recall task at the outset of the first session. All 196 images from the reproduction task were included in the memorability task, with an additional set of 196 images (foils) selected evenly from the 7 memorability score bins in the same way as the first set. All 392 images were shuffled and each image was flashed on screen for 1 second, and then subjects were given a choice to press the ‘y’ key on the keyboard to indicate that they were shown the image in the reproduction task, or the ‘n’ key to indicate that they hadn’t.

#### Analysis

For the temporal reproduction task, reproduced durations were filtered by removing all trials greater than 3 standard deviations from the mean for each subject. To examine the link between memorability and reproduced duration, and because each subject was shown the same 196 images for each of the seven possible durations, we averaged the reproduced duration for each possible image. A linear mixed model (LMM) analysis was then run on these reproduced times, with the memorability score for each image as a fixed effect and subject as a random effect. In contrast, to examine the link between reproduced duration and memorability, we performed a GLMM analysis of the binary accuracy scores for each image with the memorability score for that image and the average reproduced duration as fixed effects and subject as a random effect.

As an additional analysis, we applied a Bayesian observer-actor model to the data from the temporal reproduction task (De et al., 2021; De et al., 2023). This model, based on the work of Jazayeri and colleagues (Jazayeri & Shadlen, 2010; Remington et al., 2018) and available at https://jazlab.org/resources/, conceives of performance on a time reproduction task as arising from an initial sensory measurement (m) of the presented interval, modeled as a Gaussian distribution that scales with the size of the presented interval, that is integrated with a uniform prior distribution set to the range of presented intervals to form a posterior distribution of the interval estimate that is then corrupted by production noise (p), also modeled as a Gaussian distribution. Model parameters for the measurement and production widths were fit using Matlab’s *fminsearch* function for the reproduced durations for each of the seven memorability bins. Model fits were repeated ten times using a fitting maximum of 3000 iterations; inspection of fitted parameters indicated good convergence of results.

### Neural Network Modeling

To investigate the link between our memorability findings and computational models of vision, we implemented an artificial neural network (ANN) modeling framework, in which a comparison between behavior and network responses could be compared (Doerig et al., 2023).

For the computational model, we employed here a recurrent convolutional neural network (rCNN) of the visual system. This model, termed *BLnet* (**B**ottom-up **L**ateral Network) was designed to mimic recurrent processing within the ventral visual stream, in which individual layers project back onto themselves (Spoerer et al., 2020). Critically, this framework entails “unrolling” the model in time, such that with subsequent time steps in the model, layer-specific activity is fed back onto itself via lateral input. Thus, a single time-step refers to a single “sweep” of the model. We note that the length of the time-step here is arbitrary; indeed, the model produces identical results whether the time steps are explicitly or implicitly encoded. Rather, the model relies on a difference in the stage at which layer-specific outputs are sent via bottom-up and lateral connections (van & Kriegeskorte, 2020).

Our choice of BLnet in this case was motivated by the demonstration that rCNN models provide a better match to the sequential and time-varying nature of information flow across the ventral visual stream, as well as demonstrations that the BLnet architecture can reliably predict human reaction times and accuracy to visual stimuli (Spoerer et al., 2020; Sörensen et al., 2023). Indeed, CNNs by themselves have no access to temporal duration; by adding recurrent timesteps, even arbitrary ones, rCNNs can provide outputs that vary as a function of timstep. Here, we used the BLnet code as provided at https://github.com/cjspoerer/rcnn-sat. Here, the BLnet model was trained on object recognition using the ImageNet and Ecoset (Mehrer et al., 2021) databases across eight time steps. At the readout layer, the model provides classification output in the form of a softmax probability distribution. Crucially, this distribution is provided for each of the timesteps in the BLnet model, and as a result provides a window into how image classification is refined over time. More specifically, the softmax distribution converges on a set of image classifications over time, maximizing their probability while minimizing the probability of other categories. To quantify this, we calculated the Shannon entropy of the softmax distribution at each timestep, as done in previous work. The resulting entropy by time response thus quantifies the degree of certainty in the classification over time. By setting a response threshold on these entropy values, human responses can be predicted.

In the present study, we fed all 196 images from the memorability experiments into BLnet and calculated the resulting entropy of the eight timesteps for each one. We used the pre-trained weights for Ecoset labels for the softmax distribution, resulting in a vector of 565 category probabilities for each image; however, we also found that using ImageNet labels and weights resulted in similar findings. We then compared entropy values across the seven memorability bins to examine differences in model certainty by memorability. To compare with human performance we used the model output to set an entropy threshold for classifying images into duration categories. Specifically, we set an arbitrary threshold and categorized all images as “long” if the entropy value fell below that threshold and “short” if it fell above it. This process was repeated at each of the eight timesteps, after which the average proportion of “long” responses was calculated for each timestep. To match model performance to human responses, we repeated this process across a range of entropy scores spanning the largest to smallest entropy value in the dataset. For each threshold, as a first step, we calculated the average proportion of long responses *across* all timesteps for each of the seven memorability bins; a psychometric function was then fit to these data using the same manner as described for the bisection data of Experiment 3, from which the BP was extracted. A linear regression of the BP values across memorability bins was then fit and the slope extracted; this step was designed to mimic the “bias” effect observed in behavioral data; accordingly, a positive slope in the linear regression would indicate that the average proportion of “long” responses decreased with higher memorability bins (recall that higher bins indicate *lower* memorability scores). As a second step, we calculated the slope of a linear regression of the CV values extracted from those same psychometric functions; this step was designed to mimic the “precision” effect observed in the behavioral data; here, a positive slope would indicate that the CVs of the psychometric functions are decreasing with higher memorability bins. From these two values, we found the single entropy threshold with the lowest slope value for each effect.

## References

Minding time in an amodal representational space.. (2009). Philos Trans R Soc Lond B Biol Sci, 364, 1815–1830.

Scalar timing in memory.. (1984). Ann N Y Acad Sci, 423, 52–77.

Prospective and retrospective duration judgments: A meta-analytic review.. (1997). Psychon Bull Rev, 4, 184–197.

Temporal cognition: Connecting subjective time to perception, attention, and memory.. (2016). Psychol Bull, 142, 865–907.

Attention and the subjective expansion of time.. (2004). Percept Psychophys, 66, 1171–1189.

A theory of magnitude: common cortical metrics of time, space and quantity.. (2003). Trends Cogn Sci, 7, 483–488.

Is subjective duration a signature of coding efficiency?. (2009). Philos Trans R Soc Lond B Biol Sci, 364, 1841–1851.

Perceived time is spatial frequency dependent.. (2011). Vision Res, 51, 1232–1238.

Multiple channels of visual time perception.. (2016). Curr Opin Behav Sci, 8, 131–139.

Stimulus intensity and the perception of duration.. (2011). J Exp Psychol Hum Percept Perform, 37, 303–313.

Properties of the internal clock: first- and second-order principles of subjective time.. (2014). Annu Rev Psychol, 65, 743–771.

Life motion signals lengthen perceived temporal duration.. (2012). Proc Natl Acad Sci U S A, 109, E673–7.

Emotional modulation of interval timing and time perception.. (2016). Neurosci Biobehav Rev, 64, 403–420.

The effect of scene structure on time perception.. (2013). Q J Exp Psychol (Hove), 66, 1639–1652.

Examining visual complexity and its influence on perceived duration.. (2014). J Vis, 14, 3.

Observers exploit stochastic models of sensory change to help judge the passage of time.. (2011). Curr Biol, 21, 200–206.

Activity in perceptual classification networks as a basis for human subjective time perception.. (2019). Nat Commun, 10, 267.

Perception of duration in the parvocellular system.. (2012). Front Integr Neurosci, 6, 14.

Zwaan, R., & Zacks, J. (Eds.). (2019). Perceptual Content Not Physiological Signals, Determines Perceived Duration When Viewing Dynamic, Natural Scenes. Collabra: Psychology, 5 (1). 10.1525/collabra.234

Attentional vs computational complexity measures in observing paintings. (2009). Spatial Vision, 22 (3), 195–209. 10.1163/156856809788313138

Perceptual complexity, rather than valence or arousal accounts for distracter-induced overproductions of temporal durations.. (2014). Acta Psychol (Amst), 147, 51–59.

Encoding sensory and motor patterns as time-invariant trajectories in recurrent neural networks.. (2018). Elife, 7.

Understanding the computation of time using neural network models. (2020). Proceedings of the National Academy of Sciences, 117 (19), 10530–10540. 10.1073/pnas.1921609117

Trial-by-trial predictions of subjective time from human brain activity.. (2022). PLoS Comput Biol, 18, e1010223.

A Predictive Processing Model of Episodic Memory and Time Perception.. (2022). Neural Comput, 34, 1501–1544.

Perception and Estimation of Time. (1984). Annual Review of Psychology, 35 (1), 1–37. 10.1146/annurev.ps.35.020184.000245

Duration judgment and the experience of change.. (1983). Percept Psychophys, 33, 548–560.

Time, change, and motion: the effects of stimulus movement on temporal perception.. (1995). Percept Psychophys, 57, 105–116.

Discrete events as units of perceived time.. (2012). J Exp Psychol Hum Percept Perform, 38, 549–554.

Effects of Temporal Features and Order on the Apparent duration of a Visual Stimulus.. (2012). Front Psychol, 3, 90.

Parametric Coding of the Size and Clutter of Natural Scenes in the Human Brain.. (2015). Cereb Cortex, 25, 1792–1805.

Modeling psychophysical data at the population-level: the generalized linear mixed model.. (2012). J Vis, 12.

Understanding and Predicting Image Memorability at a Large Scale. (2015, December). 2015 IEEE International Conference on Computer Vision (ICCV). 10.1109/iccv.2015.275

Understanding Image Memorability.. (2020). Trends Cogn Sci, 24, 557–568.

Recognition memory for a rapid sequence of pictures.. (1969). J Exp Psychol, 81, 10–15.

Recognition and memory for briefly presented scenes.. (2012). Front Psychol, 3, 32.

The contributions of color to recognition memory for natural scenes.. (2002). J Exp Psychol Learn Mem Cogn, 28, 509–520.

Temporal context calibrates interval timing.. (2010). Nat Neurosci, 13, 1020–1026.

Late Bayesian inference in mental transformations.. (2018). Nat Commun, 9, 4419.

Slowing the body slows down time perception.. (2021). Elife, 10.

The role of consciously timed movements in shaping and improving auditory timing.. (2023). Proc Biol Sci, 290, 20222060.

The resiliency of image memorability: A predictor of memory separate from attention and priming.. (2020). Neuropsychologia, 141, 107408.

Predicting visual memory across images and within individuals.. (2022). Cognition, 227, 105201.

Going in circles is the way forward: the role of recurrence in visual inference.. (2020). Curr Opin Neurobiol, 65, 176–193.

Recurrent neural networks can explain flexible trading of speed and accuracy in biological vision.. (2020). PLoS Comput Biol, 16, e1008215.

Mechanisms of human dynamic object recognition revealed by sequential deep neural networks.. (2023). PLoS Comput Biol, 19, e1011169.

The parietal cortex and the representation of time, space, number and other magnitudes.. (2009). Philos Trans R Soc Lond B Biol Sci, 364, 1831–1840.

Multi-level Crowding and the Paradox of Object Recognition in Clutter.. (2018). Curr Biol, 28, R127–R133.

Dynamics of scene representations in the human brain revealed by magnetoencephalography and deep neural networks.. (2017). Neuroimage, 153, 346–358.

The spatiotemporal neural dynamics of object location representations in the human brain.. (2022). Nat Hum Behav, 6, 796–811.

Distinct ventral stream and prefrontal cortex representational dynamics during sustained conscious visual perception.. (2023). Cell Rep, 42, 112752.

Children’s use of landmarks: implications for modularity theory.. (2002). Psychol Sci, 13, 337–341.

Coding of navigational affordances in the human visual system.. (2017). Proc Natl Acad Sci U S A, 114, 4793–4798.

Large-scale dissociations between views of objects, scenes, and reachable-scale environments in visual cortex.. (2020). Proc Natl Acad Sci U S A, 117, 29354–29362.

On the (a)symmetry between the perception of time and space in large-scale environments.. (2018). Hippocampus, 28, 539–548.

Population response magnitude variation in inferotemporal cortex predicts image memorability.. (2019). Elife, 8.

Larger images are better remembered during naturalistic encoding.. (2022). Proc Natl Acad Sci U S A, 119.

Perceived image size modulates visual memory.. (2023). Psychon Bull Rev.

Blood oxygen level-dependent activation of the primary visual cortex predicts size adaptation illusion.. (2013). J Neurosci, 33, 15999–16008.

Time representations can be made from nontemporal information in the brain: an MEG study.. (2006). Cereb Cortex, 16, 1797–1808.

Priority coding in the visual system.. (2022). Nat Rev Neurosci, 23, 376–388.

Remembering the Past to See the Future.. (2021). Annu Rev Vis Sci, 7, 349–365.

The three-second subjective present: A critical review and a new proposal.. (2017). Psychol Bull, 143, 735–756.

A leaky evidence accumulation process for perceptual experience.. (2022). Trends Cogn Sci, 26, 451–461.

The timescale of perceptual evidence integration can be adapted to the environment.. (2013). Curr Biol, 23, 981–986.

Adaptive Encoding Speed in Working Memory.. (2023). Psychol Sci, 34, 822–833.

Recurrence is required to capture the representational dynamics of the human visual system.. (2019). Proc Natl Acad Sci U S A, 116, 21854–21863.

The distinct modes of vision offered by feedforward and recurrent processing.. (2000). Trends Neurosci, 23, 571–579.

Flexible timing by temporal scaling of cortical responses.. (2018). Nat Neurosci, 21, 102–110.

Controlling low-level image properties: the SHINE toolbox.. (2010). Behav Res Methods, 42, 671–684.

Human performance on the temporal bisection task.. (2010). Brain Cogn, 74, 262–272.

Painfree and accurate Bayesian estimation of psychometric functions for (potentially) overdispersed data.. (2016). Vision Res, 122, 105–123.

Different methods for reproducing time, different results.. (2014). Atten Percept Psychophys, 76, 675–681.

The neuroconnectionist research programme.. (2023). Nat Rev Neurosci, 24, 431–450.

An ecologically motivated image dataset for deep learning yields better models of human vision.. (2021). Proc Natl Acad Sci U S A, 118.

